# SNP assays for DVI: cost, time, and performance information for decision-makers

**DOI:** 10.1101/2024.05.10.593619

**Authors:** Katherine Butler Gettings, Andreas Tillmar, Kimberly Sturk-Andreaggi, Charla Marshall

## Abstract

In mass disaster events, forensic DNA laboratories may be called upon to quickly pivot their operations toward identifying bodies and reuniting remains with family members. Ideally, laboratories have considered this possibility in advance and have a plan in place. Compared with traditional short tandem repeat (STR) typing, single nucleotide polymorphisms (SNPs) may be better suited to these disaster victim identification (DVI) scenarios due to their small genomic target size, resulting in an improved success rate in degraded DNA samples. As the landscape of technology has shifted toward DNA sequencing, many forensic laboratories now have benchtop instruments available for massively parallel sequencing (MPS), facilitating this operational pivot from routine forensic STR casework to DVI SNP typing. Herein, we review the commercially available SNP sequencing assays amenable to DVI, we use data simulations to explore the potential for kinship prediction from SNP panels of varying size, and we give an example DVI scenario as context for presenting the matrix of considerations: kinship predictive potential, cost, and throughput of current SNP assay options. This information is intended to assist laboratories in choosing a SNP system for disaster preparedness.

**Highlights:** 3 to 5 bullet points (maximum 100 characters per bullet point, including spaces). Each bullet point should be a full sentence and should outline the key contributions of your manuscript and how it impacts forensic science.

- Single nucleotide polymorphisms (SNPs) are useful in disaster victim identification (DVI).
- SNP panels amenable to human identification and extended kinship are described.
- Simulations demonstrate the potential for kinship prediction from SNP panels of varying size.
- Kinship predictive potential, cost, and throughput are presented for an example DVI scenario.
- Information is intended to assist laboratories in choosing a SNP system for disaster preparedness.

## Introduction

The field of forensic genetics has recently experienced a period of rapid technological advancement with the application of single nucleotide polymorphism (SNP) markers for human DNA profiling and identification [1–6]. The increased use of SNPs has been catalyzed by the development of massively parallel sequencing (MPS) as well as forensic investigative genetic genealogy (FIGG) in support of DNA casework. SNPs represent a broad and diverse class among genomic marker types, with many forensic applications. SNP categories include autosomal, X-chromosomal, Y-chromosomal, mitochondrial DNA (mtDNA), phenotype informative, ancestry informative, and identity informative. SNPs within these categories have been variously grouped into small and large panels of markers for forensic applications. Identity informative or iiSNPs can be used for direct human identification (one-to-one matching) as well as interpreting genetic relationships between two or more individuals (in this context, sometimes referred to as kinship informative or kiSNPs). At present, there is no core set of iiSNPs for forensic DNA testing as there is for short tandem repeat (STR) loci; a core set would be needed before implementation in criminal DNA database systems such as CODIS and Interpol. Thus, the bulk of forensic DNA casework is still centered on STR typing, with iiSNPs being peripheral or complementary to traditional methods. Lacking a consensus core set of SNPs, novel forensic SNP panels have proliferated, with multiple identity and combined application (kinship/ancestry/phenotype prediction) SNP panels now in routine use.

In this review paper, we describe forensic iiSNP panels (applicable to both identity and kinship testing) and associated assays that have been published in the literature, are amenable to implementation in a typical forensic DNA laboratory and are currently available for purchase as standard products. This review does not include custom panels/assays that are not readily available, such as the QIAseq Investigator ID SNP Panel [4] and the QIAGEN MPSplex [6]. Furthermore, this review excludes population-specific SNP panels [7], large SNP panels (∼100,000 SNPs) for extended kinship applications [8], as well as SNP microarrays or whole genome sequencing (WGS) geared towards FIGG [9]. We describe the composition of the SNP panels; the overlap of markers between panels; and discuss technology, cost, and turnaround time of each panel’s associated wet lab assay(s). We further explore the potential of each panel to provide varying degrees of kinship predictions using simulations. Finally, we discuss considerations for choosing a SNP panel/assay in a DVI scenario [10]. Our aim is to provide the forensic community with a holistic examination of commercially available SNP options for DVI casework, offering a reference document for laboratories and other entities when choosing a SNP system for disaster preparedness.

## Forensic SNP Panels for Human Identification

### Early Publications of Smaller Identity SNP Panels

In the 2000’s, academic laboratories were developing small (sub-100 marker) SNP panels for human identification that could be assayed using the lower-throughput methods available at the time (e.g., SNaPshot Multiplex Kit, Applied Biosystems). Two such iiSNP panels have persisted and become absorbed into the larger assays which became possible as sequencing technologies advanced. One of these panels was developed through the efforts of the SNPforID Consortium (the SNPforID panel), which included 52 iiSNPs that were originally combined in a SNaPshot assay [11]. These SNPs were selected based on heterozygosity in three major populations (European, Asian, African) and distance between candidate markers (at least 100 kb); further, linkage disequilibrium (LD) tests for pairs of SNPs on the same chromosome demonstrated no significant deviation from expectation [11]. Subsequent publications of this panel include a forensic validation [12] and the use of the assay in parentage cases [13]. In an unrelated effort, another panel was published in 2010 as a set of 95 iiSNPs (herein called Kidd 95), with a subset of 45 iiSNPs being well spread across the chromosomes and showing “very loose or no genetic linkage with each other” among the 44 populations evaluated [14]. A related publication includes additional population data and early assay development for the subset of 45 iiSNPs, referred to hereafter as Kidd 45 [15]. Although assay performance issues have resulted in exclusion of certain SNPs, the commercially available assays described below include most of the SNPs characterized across the SNPforID and Kidd 45 panels of 52 and 45 iiSNP markers, respectively.

### Precision ID Identity Panel

The Precision ID Identity Panel (ThermoFisher Scientific, Waltham, MA), hereafter referred to as Precision ID, includes 48 iiSNPs from the SNPforID panel, 52 iiSNPs from the Kidd 95 (primarily from the Kidd 45), and 34 “upper Y-Clade SNPs” [16], for a total of 134 SNPs with an average amplicon size of 138 base pairs (bp) [17]. Other Precision ID panels may be combined with the Identity panel to reduce the cost and time of co-analyzing additional markers, such as ancestry or phenotype SNPs, Y-SNPs, and mtDNA [18]. The Identity Panel was originally released in 2014 as the HID-Ion AmpliSeq Identity Community Panel (ThermoFisher Scientific) for sequencing on the Ion Torrent Personal Genome Machine (Ion PGM) (ThermoFisher Scientific). The manufacturer currently supports sequencing on the Ion S5, S5XL or Ion GeneStudio S5 series of instruments (ThermoFisher Scientific). The difference between the three S5 configurations is analysis time; reads/output and runtime are equivalent. The Ion Chef instrument automates library preparation but increases cost and limits throughput to eight samples at a time. Manual library preparation is a less expensive option, allows for higher throughput, and uses the Ion OneTouch 2 System for amplification and enrichment. The Precision ID user manual recommends a 530 chip for sequencing; however, higher throughput methods are available from the manufacturer (e.g., 540 chip, Genexus Integrated Sequencer). While the manufacturer recommends 1 ng template DNA input, full profiles ≥ 50X coverage per marker are reported down to 50 pg template DNA input with increased amplification cycles and decreased sample multiplexing [5]. Data analysis is performed on the Ion Torrent Server with a plug-in available from ThermoFisher Scientific. The Precision ID Identity Panel primer pool has also been used in a workflow for MiSeq sequencing, with custom library preparation and bioinformatic pipeline [19,20].

### ForenSeq DNA Signature Prep Kit

The ForenSeq DNA Signature Prep Kit, hereafter referred to as Signature Prep, was originally released by Illumina in 2015 and is currently available from Verogen, which is now a part of QIAGEN (Hilden, Germany). The assay targets 94 iiSNP markers, 83 of which overlap with those in Precision ID. Additionally, 27 autosomal STR markers, 24 Y-STR markers, and 7 X-STR markers are targeted using the DNA Primer Mix A (DPMA); these co-analyzed STR markers could aid identification efforts for samples of sufficient quality. An alternative primer mix (DPMB) adds 56 ancestry and 22 phenotype SNP markers. Sequencing is performed on the MiSeq FGx instrument and output data analyzed with the Universal Analysis Software (UAS), both from Verogen/QIAGEN. The manufacturer recommends 1 ng template DNA input; however, the published developmental validation shows a 100% call rate down to 62.5 pg input DNA [21]. While the manufacturer’s procedure describes manual library preparation using a relatively simple two-step PCR process, an automated method for library preparation has been published [22].

### IDSeek OmniSNP Identity Informative SNP Typing Kit

The IDSeek OmniSNP Identity Informative SNP Typing Kit, hereafter referred to as OmniSNP, was recently released by NimaGen (Nijmegen, Netherlands). It uses reverse-complement PCR technology, which enables target enrichment and library preparation in a single PCR step [23]. The OmniSNP assay targets 85 iiSNP markers, all of which are a subset of the 94 Signature Prep iiSNPs. Seventy-nine of these 85 iiSNPs overlap with those in Precision ID; thus, 79 markers overlap across all three of these smaller iiSNP assays (Precision ID, Signature Prep, and OmniSNP). The OmniSNP libraries are designed for sequencing on a MiSeq (or MiSeq FGx in RUO mode) instrument with either a MiSeq v2 300 cycle or v3 600 cycle (or FGx) Reagent Kit. Additionally, throughput for the OmniSNP panel could be increased by using an instrument with higher output, such as a NextSeq (Illumina, San Diego, CA). A publication describing the development of this assay showed most of the SNP alleles were detected with DNA input of 60 pg [24]. The OmniSNP workflow does not have an intended bioinformatic software; however, STRait Razor Online v.0.1.7 and STRait Razor v3 were previously used for data analysis [24–26].

### FORCE

The FORCE panel was published in 2021 as an all-in-one SNP panel for forensic applications [1]. The initial panel design targeted 5,507 SNPs, including a novel kinship SNP panel of 3,935 autosomal markers, plus 140 common iiSNPs (SNPs in both Precision ID and Signature Prep), 254 ancestry SNPs, 41 phenotyping SNPs, 246 X-chromosomal SNPs, and 891 Y-chromosomal SNPs. The criteria for the kinship SNPs were designed to maximize global population diversity while minimizing genetic linkage and linkage disequilibrium and excluding clinically relevant SNPs to minimize genetic privacy concerns. Tillmar et al. demonstrated through simulations that the FORCE kinship SNPs are capable of yielding accurate relationship predictions for up to 5^th^ degree relatives with strong statistical support. The FORCE panel was first tested using an in-solution hybridization capture approach to accommodate severely degraded DNA (<75 bp fragments) [1]. In 2023, Staadig et al. evaluated the FORCE panel utilizing the QIAseq Targeted DNA chemistry (QIAGEN) and MiSeq sequencing as the basis for the laboratory workflow [27]. The QIAseq enrichment technology employs a unique primer design, which makes the method more amenable to degraded DNA than traditional PCR enrichment. Due to limitations of the enrichment approach (i.e., single primer extension), 10 SNPs were excluded from the initial panel design, resulting in a total of 5,497 SNPs in the QIAseq FORCE panel (4,073 total iiSNP and kiSNPs, all categorized herein as iiSNPs). The QIAseq FORCE method was shown to produce accurate genotype calls down to 125 pg DNA input [27]. In addition to the MiSeq (or MiSeq FGx in RUO mode), the QIAseq FORCE panel can also be sequenced on higher throughput instruments (e.g., NextSeq) [1]. Finally, an inter-laboratory study by the coauthors of this paper has shown the ability to analyze the FORCE panel using AmpliSeq technology and sequencing on an Ion platform (Ion S5/S5XL/ GeneStudio S5) (manuscript in preparation). Similar to OmniSNP, the FORCE panel does not have an intended bioinformatic software. However, QIAseq data does require specialized bioinformatic software and/or tools to facilitate the removal of PCR duplicates based upon unique molecular indexes (UMI), which are ligated to the original DNA template molecules at the beginning of library preparation. Tillmar et al. utilized CLC Genomics Workbench (QIAGEN) for data analysis and a custom script in R [28] combined with the software Merlin [29] for kinship prediction [1]. As an alternative to CLC Genomics Workbench, QIAseq FORCE data can be analyzed with a cloud-based data analysis pipeline accessible through the GeneGlobe Data Analysis Center (https://geneglobe.qiagen.com/us/analyze).

### ForenSeq Kintelligence

The ForenSeq Kintelligence kit was released by Verogen (now a part of QIAGEN) in 2021 to enable the generation of SNP profiles from forensic unknown samples for the primary purpose of searching FIGG databases while also supporting direct comparisons [3]. The 10,230 SNPs targeted for amplification by this assay are mostly geared toward kinship determination, described as having the potential to estimate 4^th^ degree relationships [30], but also include the identity, ancestry and phenotype SNP markers in Signature Prep. The laboratory procedure for Kintelligence mirrors the two-step PCR method of Signature Prep with some modification, such as an additional purification step after the first PCR. The manufacturer recommends 1 ng template DNA input, yet recent publications have shown >85% call rates for DNA inputs as low as 50 pg to 0.1 ng input [3,31]. The high level of marker multiplexing limits the manufacturer’s recommended throughput to three libraries in each MiSeq FGx sequencing run for FIGG purposes. A published internal validation evaluated sequencing pools of two to four sample libraries for FIGG purposes, depending on sample quality [32]. The manufacturer’s software (Universal Analysis Software or UAS) facilitates long-range kinship analysis in GEDmatch PRO, and this search is included in the cost of the library preparation kit.

Of particular relevance to this review, a recent publication from the manufacturer describes the use of a higher throughput workflow for DVI scenarios that allows for lower call rates [33]. This is possible because direct kinship testing for DVI can be performed with fewer SNPs than are needed for a GEDmatch PRO search. A library preparation kit for this new, higher throughput workflow has recently been released as the ForenSeq Kintelligence HT kit, with a manufacturer’s recommended limit of 12 post-mortem (PM) or 36 ante-mortem (AM) samples per MiSeq FGx run. The Kintelligence HT kit contains the same 10,230 SNPs as the standard Kintelligence kit but is a less expensive option for two reasons: 1) the library preparation kit has a decreased cost per sample compared to the original Kintelligence library preparation kit, and 2) increasing the number of samples per sequencing run decreases the per sample sequencing cost. Additionally, the purchase of the Kintelligence HT library preparation kit includes access to a local UAS version geared specifically toward DVI database matching (personal communication with QIAGEN).

## Cross-Panel SNP Overlap

All SNPs included in these five commercially available panels, along with SNP categories and GRCh38 coordinates, are listed in Supplemental File S1. These SNP panels vary greatly in the number of iiSNPs they contain, but each has some overlap with other panels, as shown in Table 1. The levels of overlap between the differently sized panels (Kintelligence, FORCE and Signature Prep, as an exemplar small panel) are shown in Figure 1. Notably, there is significant overlap between the smallest three identity SNP panels: Signature Prep, Precision ID, and OmniSNP, as shown in Supplemental File S2. This high proportion of overlapping markers (>84% for each pairwise comparison) owes to the panels’ designs primarily incorporating iiSNPs from the same two early panels (SNPforID and Kidd 45), as previously described. Additionally, nearly all of the SNP markers in these small panels are contained within the larger FORCE and Kintelligence SNP panels, by design. Minor variations exist (including or excluding slightly different markers) due to varying chemistries and compatibilities, leaving 76 iiSNPs in common between all five panels.

**Figure 1.**
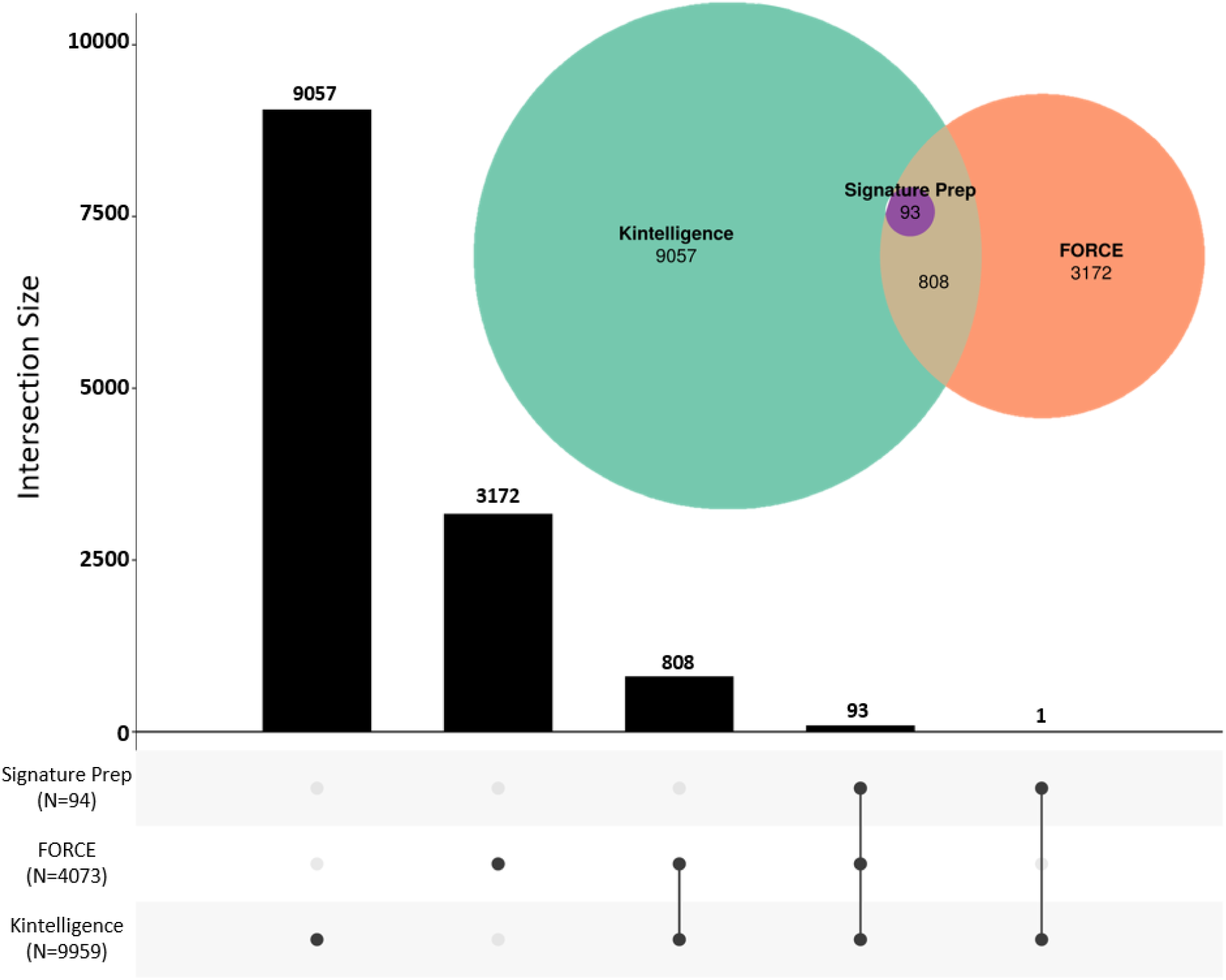
Area-proportional Euler plot (colored circles) overlaid on an Upset plot (black and white), showing overlap between the iiSNPs implemented in Signature Prep, QIAseq FORCE, and Kintelligence. Signature Prep was included as an exemplary small panel to represent the overlap with QIAseq FORCE and Kintelligence. All pairwise overlaps including those with the Precision ID and OmniSNP panels are shown in Table 1. Plots generated with R packages eulerr and UpSetR (https://shiny2.imetalab.ca/shiny/rstudio/SetExplorer/).

**Table 1.**
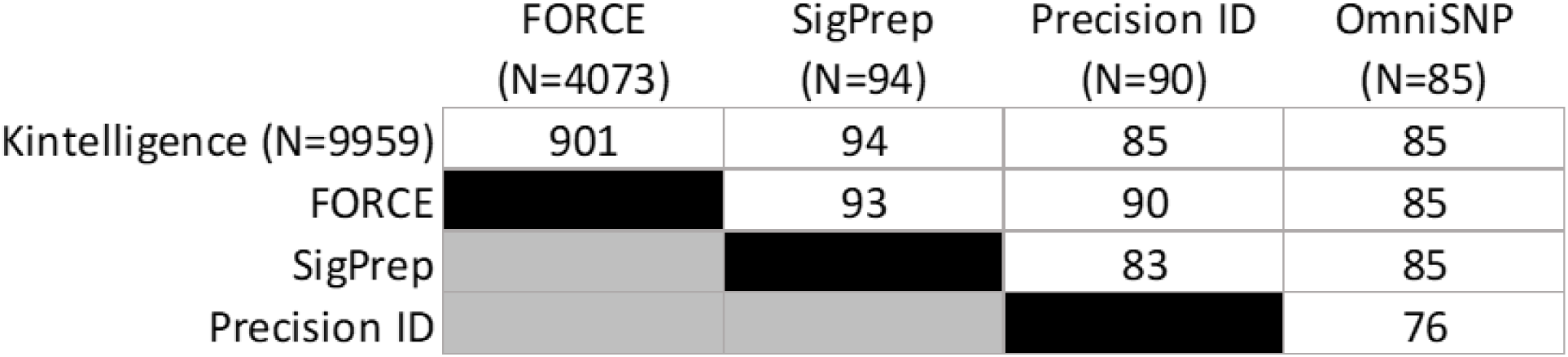
Overlap between the iiSNPs implemented in Kintelligence, QIAseq FORCE, Signature Prep, Precision ID, and OmniSNP.

Beyond the 76 iiSNPs in common between all five panels, QIAseq FORCE and Kintelligence contain 826 additional overlapping identity markers, for a total of 22% overlap between these two larger panels (with respect to the number of total FORCE SNPs). While the criteria for selecting identity markers were similar for both FORCE and Kintelligence, this relatively low level of overlapping markers speaks to the abundance of iiSNPs in the human genome.

## Kinship Prediction

The possibility for kinship prediction from both full and partial evidentiary/unknown profiles was assessed through data simulations utilizing each of the five SNP panels. Full profiles were assumed for all reference/known samples for all conditions, and partial profiles were generated for the evidentiary/unknown profiles comprising a random number of markers ranging from 30% to 100% of a full profile for each simulation event. In brief, the simulations were performed using the software Merlin [29]. Merlin was used both to create the pedigree-based DNA data and to calculate likelihoods for the tested hypotheses. European allele frequencies from the 1000 Genomes dataset [34] were used, and genetic position information from the Rutgers map was used to account for genetic linkage [35]. For each simulated kinship case scenario, the likelihood ratio (LR) was calculated as LR = Pr (DNA|H1)/Pr (DNA|H2), where H1 was the hypothesis representing the related case scenario, and H2 was the hypothesis representing the alternative, in which the tested individuals are unrelated. The tested relationships include the following: 1^st^ – parent/child, full sibling; 2^nd^ – half-sibling, aunt/uncle, grandchild, etc.; 3^rd^ – first cousin, great-grandchild, etc.; 4^th^ – first cousin once removed (1C1R), etc.; and 5^th^ degree – second cousin, etc. For each case scenario and true hypothesis, 10,000 simulation events were performed. A front-end script written in R [28] was used to format and manage input files, and R was also used to handle the outputs from Merlin and for plotting.

Figure 2 shows box plots depicting log10 LRs comparing each of five relationship categories to unrelated for Signature Prep (as an exemplary small panel), QIAseq FORCE, and Kintelligence (using iiSNPs only for each panel). Median log10 LRs for 1^st^ degree relationship predictions from full evidentiary/known profiles were: 6 for Signature Prep, 447 for FORCE, and 1300 for Kintelligence. However, when considering partial profiles, each of the median log10 LRs for 1^st^ degree relationship predictions was reduced by approximately 40%, to 3.5 for Signature Prep, 272 for FORCE, and 784 for Kintelligence. Results from all five panels (OmniSNP and Precision ID alongside the data in Figure 2) are shown in Supplemental File S3. Across all panels, the median log10 LRs generally decreased by a factor of three for every single-step increase in relationship distance. For example, full profile median log10 LRs for Kintelligence were 1300 for 1^st^ degree, 300 for 2^nd^ degree, 120 for 3^rd^ degree, 50 for 4^th^ degree, and 20 for 5^th^ degree. In general, the higher the SNP count and the closer the relationship, the higher the log10 LRs that were produced via simulations.

**Figure 2.**
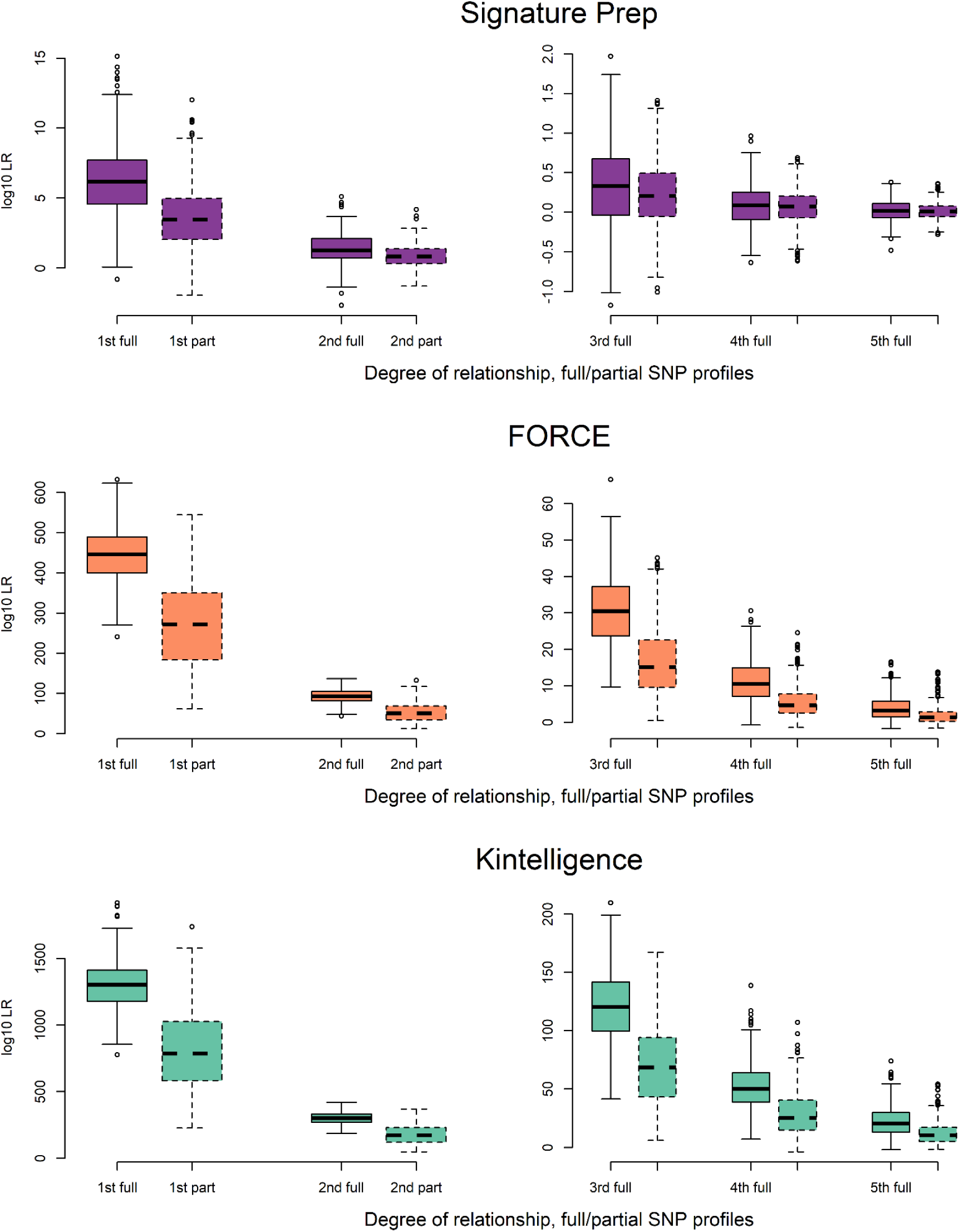
Box plots depicting log10 LRs comparing each of five relationship categories to unrelated for Signature Prep, FORCE, and Kintelligence. Signature Prep is included as an exemplary small panel; results for all five panels can be found in Supplementary File S3. Full iiSNP profile results are shown with solid black outline, and partial profile results are shown with the dashed line. The tested relationships include the following: 1^st^ – parent/child, full sibling; 2^nd^ – half-sibling, aunt/uncle, grandchild, etc.; 3^rd^ – first cousin, great-grandchild, etc.; 4^th^ – first cousin once removed (1C1R), etc.; and 5^th^ degree – second cousin (2C), etc.

Figure 3 shows exceedance plots for Signature Prep (as an exemplary small panel), FORCE, and Kintelligence to visualize the proportion of relationships predicted at increasing log10 LR thresholds. Exceedance plots for all five panels by relationship (OmniSNP and Precision ID alongside the data shown in Figure 3) are shown in Supplemental File S4. Exceedance plots by panel for full and partial profiles for all five SNP panels are shown in Supplemental File S5 and the underlying data is given in table format in Supplemental File S6. The log10 LR thresholds correspond to forensic verbal equivalency scales ranging from 0 (LR=1, *uninformative*), 1 (LR=10, *limited support*), 2 (LR=100, *moderate support*), 3 (LR=1,000, also *moderate support*), 4 (LR=10,000, *strong support*), 5 (LR=100,000, also *strong support*), and 6 (LR=1,000,000, *very strong support*) [36]. The Signature Prep log10 LRs indicate that 82% of 1^st^ degree and 0.8% of 2^nd^ degree relationships were predicted with *strong* or *very strong support*, while 3^rd^ degree relationships and beyond did not achieve a verbal equivalent of *strong support*. The QIAseq FORCE panel produced log10 LRs corresponding to *very strong support* for all 1^st^ through 3^rd^ degree relationships, while 91% of 4^th^ degree and 44% of 5^th^ degree relationships exceeded the *strong support* log10 LR threshold. All Kintelligence 1^st^ to 4^th^ degree relationship predictions yielded *very strong support*, and 95% of 5^th^ degree relationships were predicted with *strong* or *very strong support* for this large SNP panel. These log10 LR values and corresponding verbal equivalents assume complete coverage of all iiSNPs, which is less likely with the Kintelligence HT workflow’s maximum manufacturer recommended number of samples in a sequencing run (12 PM or 36 AM samples). Based on the results published in [33], the simulation results of the partial profiles in Supplemental Files S5 and S6 better represent expectations with Kintelligence HT. Regardless, for either full or partial Kintelligence profiles, differences in the ability to achieve the log10 LR verbal equivalent of *strong support* were only found for 5^th^ degree relationships (95% for full and 81% for partial profiles).

**Figure 3.**
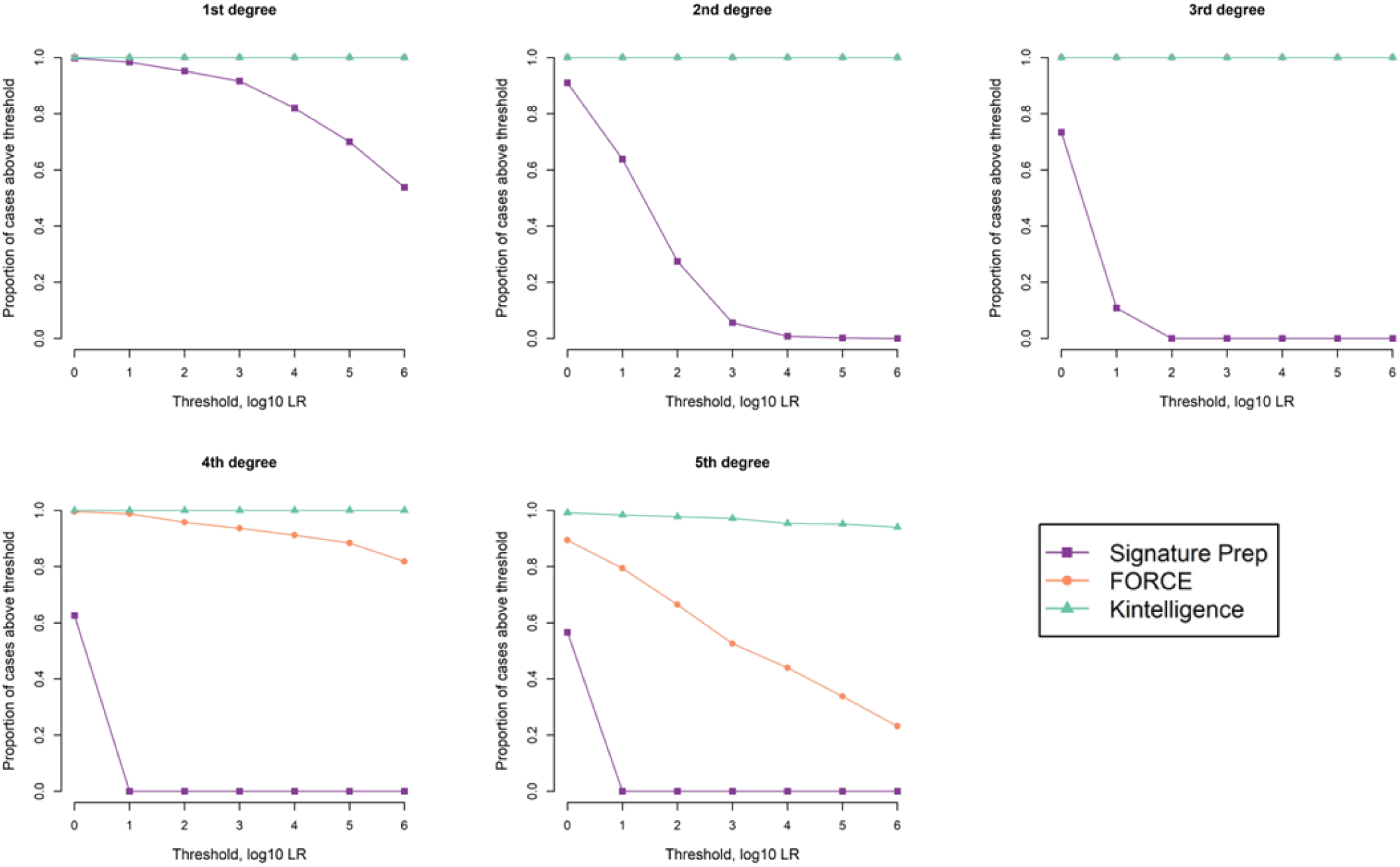
Exceedance plots by relationship category indicating the proportion of cases exceeding the following log10 LR thresholds with corresponding forensic verbal scale equivalencies: 0 (LR=1, *uninformative*), 1 (LR=10, *limited support*), 2 (LR=100, *moderate support*), 3 (LR=1,000, also *moderate support*), 4 (LR=10,000, *strong support*), 5 (LR=100,000, also *strong support*), and 6 (LR=1,000,000, *very strong support*). The results are shown for Signature Prep (as an exemplary small SNP panel), QIAseq FORCE, and Kintelligence. Additional data can be found in Supplemental Files: S4 contains exceedance plots by relationship category for all five panels, S5 contains exceedance plots by panel for all five relationship categories and full/partial profiles, and S6 contains these probabilities in table format.

Even though all simulations were conducted under the assumption of relatedness, events with LR<1 (log10 LR<0) were also obtained, as can be observed in Figure 2 box plots. Moreover, the proportion of log10 LR>0 values in the Figure 3 exceedance plots is not always 1. Since the LR in these cases has been calculated to be <1, this provides support against a relationship, which may lead to false exclusions. Therefore, it is important to conduct simulation studies like this for the degrees of relationship expected to be tested in the laboratory to understand the frequency of possible false exclusions based on the chosen SNP panel.

## Cost-Time Assessment

Cost, time, and performance are parameters that often impact the decision to implement a new technology within a laboratory. Though capital expenditures such as instrumentation and analysis software prices are primary cost factors in the decision-making process, per sample cost and processing time are key considerations in the selection of a SNP panel best suited for DVI scenarios as well as other forensic applications. To assist decision makers in forensic laboratories that may be considering implementation of a SNP panel for DVI and other forensic applications, cost and time estimates were determined for Precision ID, Signature Prep, OmniSNP, QIAseq FORCE, and Kintelligence HT.

The per sample cost focused solely on kit reagents required for manual processing (i.e., library preparation and enrichment reagents, indexes, purifying beads) and is based on list prices in the U.S. at the time of writing (May 2024). Consumables and quantitation reagents were not included in the cost as these items would be relatively similar between methods. Sequencing cost was incorporated into the per sample cost based upon the maximum batch size (i.e., maximum number of libraries that can be multiplexed in a single sequencing run) recommended by the manufacturer and/or based upon published data for specific sample types given a particular sequencing approach (Table 2). AM samples are typically high-quality reference-type samples such as buccal swabs from family members used to identify missing individuals. These samples can be processed in relatively large batches (e.g., 96 samples) since they typically contain high quantities of intact DNA without the presence of inhibitors (e.g., bacterial DNA). The batch size would be reduced as needed for PM samples such as bones and tissue, depending on DNA quantity and quality (degree of fragmentation) and environmental contaminants specific to the DVI incident (e.g., soil, saltwater, etc.). Alternatively, higher throughput sequencing methods may be employed, which would increase the batch size and thus reduce the sequencing component of per sample cost (Supplemental File S7). These cost estimates are based on the standard workflows recommended by the manufacturers and/or in published literature, performed manually and using the equipment most likely to be available within a forensic laboratory at this time. Additional modifications to these methods may reduce the per sample cost.

**Table 2.** Estimates of cost and time for processing with five commercial SNP panels. The batch size for ante-mortem (AM) and post-mortem (PM) samples are based on manufacturer protocols or previously published studies with the specified sequencing approach. Cost estimates, shown in U.S. dollars (USD), include only sample preparation and sequencing reagents. Time estimates are based on manual processing by a single scientist for a batch of samples (either AM or PM). The sample preparation time listed below includes the hands-on time (noted in the parentheses) as well as time on an instrument for PCR, incubations, and quantitation steps. Sequencing time includes post-run data processing as well as templating on the Ion Chef for the Precision ID workflow.

| SNP Panel | Sequencing Approach<br>(Instrument / Kit) | Batch Size |  | Cost in USD per Sample |  |  | Batch Processing Time (Hours) |  |  |
| --- | --- | --- | --- | --- | --- | --- | --- | --- | --- |
|  |  | AM | PM | Sample<br>Preparation<br>Only | Total (including<br>Sequencing) |  | Sample<br>Preparation<br>(Hands-On) | Sequencing<br>and Post-Processing† | Total |
|  |  |  |  |  | AM | PM |  |  |  |
| Precision ID | Ion S5 /<br>530 chip | 96 | 32 | 165 | 175 | 195 | 11.5 (6.0) | 17.0 | 28.5 |
| Signature Prep | MiSeq FGx /<br>FGx Reagent Kit | 96 | 32 | 60 | 85 | 125 | 7.5 (3.5) | 30.0 | 37.5 |
| OmniSNP | MiSeq /<br>MiSeq V2 300 Cycle Kit | 96 | 32 | 30 | 45 | 75 | 10.0 (3.0) | 26.0 | 36.0 |
| QIAseq FORCE | MiSeq FGx /<br>FGx Reagent Kit | 48 | 16 | 155 | 200 | 285 | 12.5 (7.5) | 30.0 | 42.5 |
| Kintelligence HT | MiSeq FGx /<br>FGx Reagent Kit | 36 | 12 | 925 | 985 | 1100 | 9.0 (3.5) | 30.0 | 39.0 |

As shown in Table 2, the majority of the cost per sample is allocated for library preparation and enrichment reagents, averaging 71% across all five panels; therefore, differences in batch size have a relatively small effect on total cost. Overall, the OmniSNP panel has the lowest estimated cost per sample, followed by Signature Prep. However, the latter includes STRs and (optional) ancestry and phenotype SNPs, which would decrease the per marker cost. Precision ID has the highest estimated cost per sample of the small SNP panels due to the higher cost of sample preparation reagents. For the small SNP panels, sequencing adds very little cost per sample due to the large batch sizes (96 AM and 32 PM samples). In contrast, the smaller batch sizes for QIAseq FORCE (48 AM and 16 PM samples) and Kintelligence HT (36 AM and 12 PM samples) result in higher sequencing costs per sample. Notably, QIAseq FORCE sample preparation cost is lower than Precision ID, despite having substantially more SNPs, but the reduced batch size increases the total per sample cost (Table 2). The QIAseq FORCE and OmniSNP per sample cost can be reduced with the use of a higher throughput sequencing platform such as the NextSeq 550 (Supplemental File S7), unlike Signature Prep and Kintelligence HT that were specifically designed to be sequenced on a MiSeq FGx instrument. While Kintelligence HT has the highest cost per sample by far of the workflows shown in Table 2, it is important to consider that the cost of the Kintelligence HT kit incorporates the price of the UAS Kintelligence HT analysis module for direct kinship comparisons. Additionally, the original Kintelligence workflow has a higher cost per sample than Kintelligence HT (Supplemental File S7); this smaller Kintelligence batch size is optimized for FIGG investigations and includes the fee to search GEDmatch Pro.

As with the cost estimates, the standard workflow recommended by the manufacturer and most likely sequencing approach were used to estimate the SNP panel batch processing times. These time estimates are summarized in Table 2 and broken down further in Supplemental File S8. Processing time estimates assume manual library preparation by a single scientist. Hands-on and instrument times were estimated based on manufacturer guidance (i.e., user manuals or promotional material) and/or user experience. When comparing processing times, it is important to take batch size into account as certain methods are more amenable to high-throughput processing or have greater multiplexing options available (Supplementary File S7). Automation may reduce processing time, particularly hands-on time, but reagent volume overages required for automated pipetting and other limitations may increase cost. For example, the Ion Chef can perform enrichment and library preparation for the Precision ID workflow; however, the Ion AmpliSeq Kit for Chef DL8 reduces multiplexing capabilities due to the number of indexes included in the automated kit and thereby increases per sample cost by approximately 30% (Supplementary File S7). Before deciding to implement an automated method, a cost-benefit analysis should evaluate time to implement versus time saved and other factors (e.g., reduction in processing errors, reproducibility).

When considering time from extracted DNA to SNP genotypes, the shortest processing time was estimated for Precision ID, taking just over one day for a batch of samples (Table 2). OmniSNP, Signature Prep and Kintelligence HT have similar processing times (36-39 hours), including about 3 hours of hands-on time required by all three kits. In contrast, QIAseq FORCE has the longest processing time, taking approximately 42 hours for a sample batch (Table 2). This is primarily due to the more labor-intensive sample preparation, which is estimated to require 12.5 hours to complete and has numerous hands-on steps (as shown in Supplemental File S8). Kintelligence HT and QIAseq FORCE both have reduced batch sizes compared to the three smaller panels thus, sequencing larger numbers of samples (e.g., 96) with Kintelligence HT or QIAseq FORCE requires multiple runs, increasing the overall time. Sample-to-genotype time is crucial in most DVI scenarios, therefore processing time and throughput (i.e., batch size) are critical parameters to consider when comparing SNP panels for this application. The total time required for sample-to-identification also includes relationship matching, statistical calculations and reporting. These final steps are beyond the scope of this review because most of these workflows do not have a manufacturer recommended software tool for relationship testing and, in any case, the time required would depend largely on the number/types of samples, available references, and user expertise.

## Application to a DVI Scenario

Precision ID, OmniSNP, Signature Prep, QIAseq FORCE, and Kintelligence HT were assessed based on the cost, time and performance in the context of an example historical DVI scenario. Kenya Air Flight 507, with 114 passengers and crew members, crashed shortly after takeoff from Douala International Airport (Cameroon) in 2007. The identification effort was undertaken by the International Commission for Missing Persons (ICMP), which included STR DNA testing of 316 PM samples from the crash site and 114 donor reference (AM) samples [37]. The cost and time to process these 430 DVI samples with SNP assays described in this review were based on the estimates presented in Table 2. In addition to the parameters applied above to determine the estimates (e.g., batch sizes for AM and PM samples, manual processing by a single scientist, etc.), it was assumed that only one sequencing instrument was available in the laboratory for this DVI incident. The kinship predictive potential of each SNP panel was used as the measure of performance. In most DVI incidents, the most common relationships between the decedents (PM samples) and the submitted references (AM samples) are 1^st^ degree (i.e., parent-offspring, full siblings). Therefore, the performance parameter used in this assessment was the proportion of 1^st^ degree relationship simulations that were able to achieve *strong* statistical support (LR ≥ 10,000) as shown in Figure 3 and Supplemental File S4.

The cost and time estimates for these five methods as applied to the DVI scenario of the Kenya Airlines Flight 507 crash are shown in Supplemental File S9. Cost, time, and performance of Signature Prep (as an exemplary small panel), QIAseq FORCE, and Kintelligence HT in this DVI scenario are shown in Figure 4. Based on these estimates, Signature Prep would be the lowest cost and fastest option to process the 430 total samples. With increasing numbers of SNPs, QIAseq FORCE would cost more than double that of Signature Prep and take almost twice as long to process the samples. Kintelligence HT would be the most expensive option and would take the longest to process the AM and PM samples (Supplemental File S9). QIAseq FORCE and Kintelligence HT would be expected to perform similarly in terms of the kinship prediction potential for 1^st^ degree relationships (for both, 100% of the simulations achieved log10 LRs with a verbal equivalent of *strong support*), whereas the Signature Prep iiSNPs have a slightly reduced kinship prediction potential for 1^st^ degree relationships (Supplemental File S6). However, the performance of Signature Prep would likely be sufficient in the context of many DVI scenarios involving primarily 1^st^ degree relationships, and the included STRs may provide additional information if the PM sample quality is sufficient. Though the smaller panels (Signature Prep, Precision ID, and OmniSNP) perform similarly in terms of iiSNP kinship prediction potential (Supplemental File S10), cost and time estimates within a DVI scenario differentiate these three SNP panels. As shown in Supplemental Files S9 and S10, Precision ID would have the shortest processing time for this DVI scenario (10 days) but the highest cost of the smaller SNP panels (about $80,000). The OmniSNP panel has a substantially lower cost (less than $29,000) compared to the other kits, with a relatively short processing time of 14 days.

**Figure 4.**
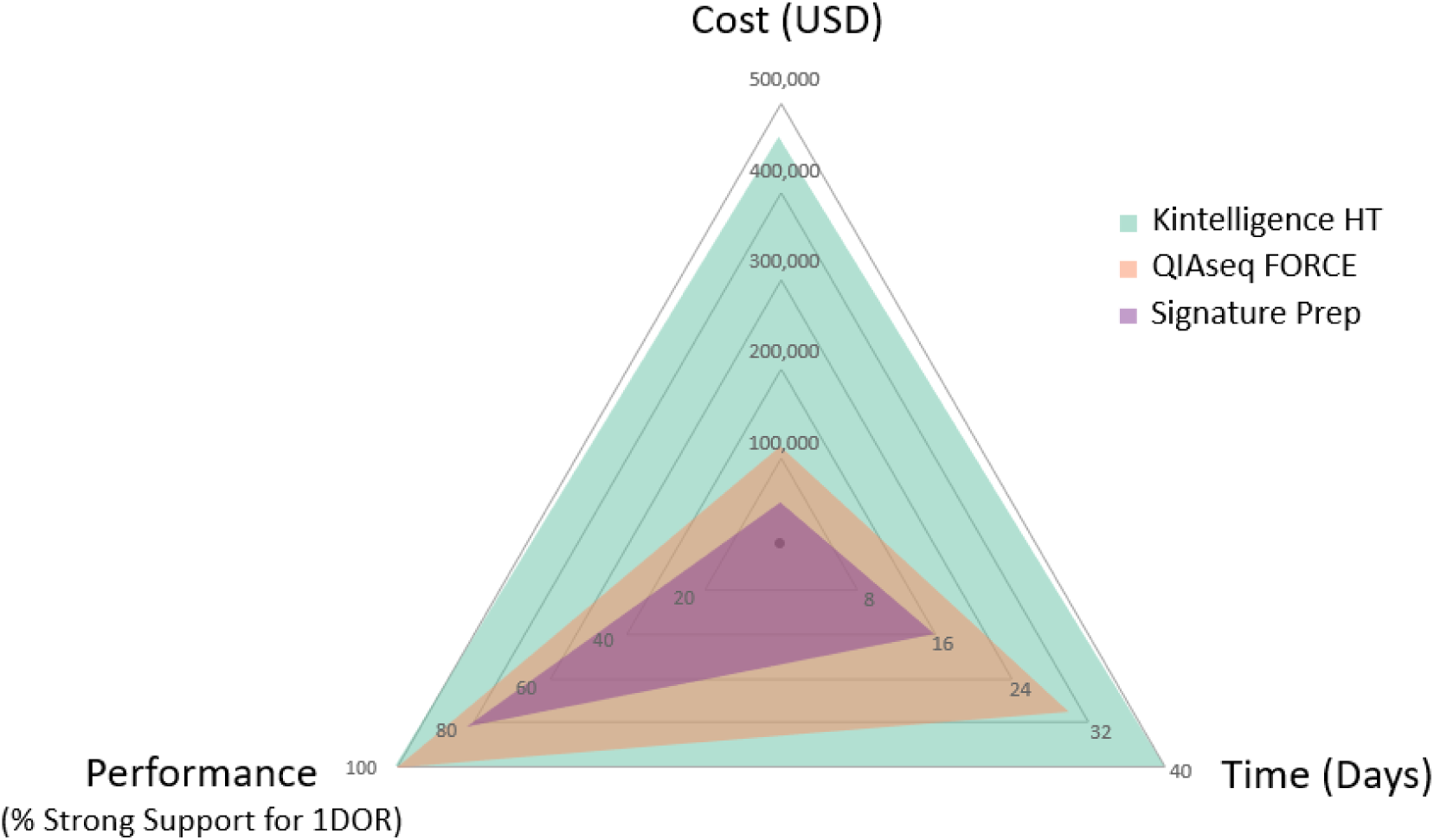
Radar plot scaling the cost, time, and performance of Signature Prep (as an exemplar small panel, purple), QIAseq FORCE (orange), and Kintelligence HT (green) in the context of an example DVI scenario (Kenya Airlines Flight 507 crash in 2007). Cost and time are the total amounts required to process all 430 samples given the batch sizes established for each sample type (ante-mortem and post-mortem). Performance is shown as the proportion of simulated kinship predictions with *strong* statistical support (LR ≥ 10,000) simulated for 1^st^ degree relatives (1DOR, i.e., parent-child, full siblings).

Though the cost and processing time of the smaller SNP panels is less (Supplemental File S9), a larger SNP panel like Kintelligence HT or QIAseq FORCE would be needed if more distant relationships were involved (i.e., grandparents, aunts/uncles, cousins) since there is a substantial loss in kinship prediction potential for the panels with fewer SNPs (Supplemental Files S5 and S6). Differentiation in performance between Kintelligence HT and QIAseq FORCE occurs around 4^th^ degree relationships and is more pronounced with 5^th^ degree relationships.

Thus, AM sample availability may need to be considered when choosing a method for particular DVI scenarios, such as historical incidents in which AM samples from close family members are not available. PM sample quality is an additional consideration for specific scenarios. While amplification-based approaches (including all five methods considered for this cost-time assessment) are expected to perform similarly, other methods (such as the capture hybridization implementation of the FORCE panel) may be more robust to degraded DNA and better suited to historical DVI incidents.

## Conclusion

While forensic scientists and laboratory directors may already be evaluating SNP methods for forensic casework, larger entities such as state and national laboratories may be considering or developing SNP typing capabilities for mass disaster event preparedness. Ideally, a SNP solution implemented for routine casework could also be applicable to DVI scenarios, making it easier to shift the focus of laboratory operations when needed. Additionally, if the laboratory has implemented a smaller SNP panel for routine casework, this less-expensive option could serve as the laboratory’s primary approach for identity matching and relationship prediction with 1^st^ degree relatives in a DVI scenario, and then a larger SNP panel could be used (sample quantity permitting) for more distant relationship testing or long-range familial searching of e.g., genetic genealogy databases.

There are numerous factors that can impact the selection and use of a SNP panel in a DVI scenario. Sequencing platforms and data analysis software are often the primary factors considered for the implementation of new methods in a forensic laboratory, as these are largest, upfront capital expenditures. However, the factors described here--cost/time of sample processing and performance in kinship predictions--may better inform the selection of a SNP panel given the DVI scenarios likely to be encountered by a laboratory, depending on its mission and scope of forensic identification services. This review represents a snapshot in time at a pivotal point in technology development for SNP-based human identification MPS assays, which may help inform these important decisions.

## Supporting information

Supplemental Files S1 through S10

## Acknowledgements

The authors thank Keith Elliott and Laurence Devesse (QIAGEN), Pieter van Oers (NimaGen), and Rob Lagace (ThermoFisher), as well as Becky Steffen and Kevin Kiesler (NIST), Adam Staadig (NBFM), and Daniele Podini (GWU) for assay information and discussions.

## Disclaimer

AFMES-AFDIL: The assertions herein are those of the authors and do not necessarily represent the official position of the United States Department of Defense, the Defense Health Agency, or its entities including the Armed Forces Medical Examiner System. Any mention of commercial products was done for scientific transparency and should not be viewed as an endorsement of the product or manufacturer.

NIST: Points of view are those of the authors and do not necessarily represent the official position or policies of the U.S. Department of Commerce. Certain commercial software, instruments, and materials are identified in order to specify experimental procedures as completely as possible. In no case does such identification imply a recommendation or endorsement by NIST, nor does it imply that any of the materials, instruments, or equipment identified are necessarily the best available for the purpose.

## Supplemental Information

*Supplemental File S1*. All SNPs included in all five panels, including indicated assay options (FORCE QIAseq and FORCE myBaits; Signature Prep DPMA and DPMB). Category “Identity” includes both Kinship and Identity SNPs, sometimes differentiated by the developer/manufacturer.

*Supplemental File S2*. Area-proportional Euler plot (colored circles) overlaid on an Upset plot (black and white), showing overlap between the iiSNPs implemented in ForenSeq DNA Signature Prep Kit, Precision ID Identity Panel, and ID Seek OmniSNP Identity Informative SNP Typing Kit.

*Supplemental File S3*. Box plots depicting log10 LRs comparing each of five relationship categories to unrelated for OmniSNP, Precision ID, Signature Prep, FORCE, and Kintelligence. Full profile results are shown with solid black outline, and partial profile results are shown with the dashed line. The tested relationships include the following: 1st – parent/child, full sibling; 2nd – half-sibling, aunt/uncle, grandchild, etc.; 3rd – first cousin, great-grandchild, etc.; 4th – first cousin once removed (1C1R), etc.; and 5th degree – second cousin (2C), etc.

*Supplemental File S4*. Exceedance plots indicating the proportion of cases exceeding the following log10 LR thresholds with corresponding forensic verbal scale equivalencies: 0 (LR=1, uninformative), 1 (LR=10, limited support), 2 (LR=100, moderate support), 3 (LR=1,000, also moderate support), 4 (LR=10,000, strong support), 5 (LR=100,000, also strong support), and 6 (LR=1,000,000, very strong support). Results are shown by relationship category (1st through 5th degree) for OmniSNP, Precision ID, Signature Prep, QIAseq FORCE, and Kintelligence iiSNPs.

*Supplemental File S5*. Exceedance plots indicating the proportion of cases exceeding the following log10 LR thresholds with corresponding forensic verbal scale equivalencies: 0 (LR=1, uninformative), 1 (LR=10, limited support), 2 (LR=100, moderate support), 3 (LR=1,000, also moderate support), 4 (LR=10,000, strong support), 5 (LR=100,000, also strong support), and 6 (LR=1,000,000, very strong support). Full and partial profile results for 1st through 5th degree relationship categories are shown by iiSNP panel for OmniSNP, Precision ID, Signature Prep, QIAseq FORCE, and Kintelligence.

*Supplemental File S6*. Probabilities of cases exceeding the following log10 LR thresholds with corresponding forensic verbal scale equivalencies: 0 (LR=1, uninformative), 1 (LR=10, limited support), 2 (LR=100, moderate support), 3 (LR=1,000, also moderate support), 4 (LR=10,000, strong support), 5 (LR=100,000, also strong support), and 6 (LR=1,000,000, very strong support). Full and partial profile results for 1st through 5th degree relationship categories are shown by iiSNP panel for OmniSNP, Precision ID, Signature Prep, QIAseq FORCE, and Kintelligence.

*Supplemental File S7*. Estimates of cost and time for processing with five commercial SNP panels, including the approach used for the cost-time assessment (no shading, also shown in Table 2), higher throughput methods (green shading), and an automated method (blue shading). The batch size for antemortem (AM) and post-mortem (PM) samples are based on manufacturer protocols or previously published studies with the specified sequencing approach. Cost estimates include only sample preparation and sequencing reagents. Time estimates are based on manual processing by a single scientist for a batch of samples (either AM or PM). The sample preparation time listed below includes the hands-on time (noted in the parentheses) as well as time on an instrument for PCR, incubations, and quantitation steps. Sequencing time includes post-run data processing as well as templating on the Ion Chef for the Precision ID workflow.

*Supplemental File S8*. Workflow Gantt charts for all five assays based on manufacturer recommendations for antemortem samples. Hands-on time = orange, hands-off time in library preparation = dark blue, Ion Chef templating = light blue, sequencing and data processing = green.

*Supplemental File S9*. Cost and time estimates of five commercial SNP panels given the batch sizes established for each sample type (AM and PM), in the context of an example DVI scenario (Kenya Airlines Flight 507 crash in 2007, 430 total samples).

*Supplemental File S10*. Radar plot scaling the cost, time, and performance of Signature Prep, Precision ID, and OmniSNP (blue) in the context of an example DVI scenario (Kenya Airlines Flight 507 crash in 2007). Cost and time are the total amounts required to process all 430 samples given the batch sizes established for each sample type (AM and PM). Performance is shown as the proportion of simulated kinship predictions with strong statistical support (LR ≥ 10,000) simulated for 1st degree relatives (1DOR, i.e., parent-child, full siblings).

## References

[1] A. Tillmar, K. Sturk-Andreaggi, J. Daniels-Higginbotham, J.T. Thomas, C. Marshall, The FORCE Panel: An all-in-one SNP marker set for confirming investigative genetic genealogy leads and for general forensic applications, Genes. 12 (2021) 1968.

[2] J. Ruiz-Ramírez, M. de la Puente, C. Xavier, A. Ambroa-Conde, J. Álvarez-Dios, A. Freire-Aradas, et al., Development and evaluations of the ancestry informative markers of the VISAGE Enhanced Tool for Appearance and Ancestry, Forensic Science International: Genetics. 64 (2023) 102853.

[3] J. Antunes, P. Walichiewicz, E. Forouzmand, R. Barta, M. Didier, Y. Han, et al., Developmental Validation of the ForenSeq® Kintelligence Kit, MiSeq Fgx® Sequencing System and ForenSeq Universal Analysis Software, Forensic Science International: Genetics. (2024) 103055.

[4] I. Grandell, R. Samara, A.O. Tillmar, A SNP panel for identity and kinship testing using massive parallel sequencing, Int. Journal of Legal Med. 130 (2016) 905–914.

[5] C. Turchi, C. Previderè, C. Bini, E. Carnevali, P. Grignani, A. Manfredi, et al., Assessment of the Precision ID Identity Panel kit on challenging forensic samples, Forensic Science International: Genetics. 49 (2020) 102400.

[6] C. Phillips, J. Amigo, A.O. Tillmar, M.A. Peck, M.d.l. Puente, J. Ruiz-Ramírez, et al., A compilation of tri-allelic SNPs from 1000 Genomes and use of the most polymorphic loci for a large-scale human identification panel, Forensic Science International: Genetics. 46 (2020) 102232.

[7] S. Zhang, Y. Bian, A. Chen, H. Zheng, Y. Gao, Y. Hou, et al., Developmental validation of a custom panel including 273 SNPs for forensic application using Ion Torrent PGM, Forensic Science International: Genetics. 27 (2017) 50–57.

[8] E.M. Gorden, E.M. Greytak, K. Sturk-Andreaggi, J. Cady, T.P. McMahon, S. Armentrout, et al., Extended kinship analysis of historical remains using SNP capture, Forensic Science International: Genetics. 57 (2022) 102636.

[9] E. Greytak, S. Wyatt, J. Cady, C. Moore, S. Armentrout, Investigative genetic genealogy for human remains identification, Journal of Forensic Sciences (2024).

[10] M. Prinz, A. Carracedo, W.R. Mayr, N. Morling, T.J. Parsons, A. Sajantila, et al., DNA Commission of the International Society for Forensic Genetics (ISFG): recommendations regarding the role of forensic genetics for disaster victim identification (DVI), Forensic Science International: Genetics. 1 (2007) 3–12.

[11] J.J. Sanchez, C. Phillips, C. Borsting, K. Balogh, M. Bogus, M. Fondevila, et al., A multiplex assay with 52 single nucleotide polymorphisms for human identification, Electrophoresis. 27 (2006) 1713–1724.

[12] E. Musgrave-Brown, D. Ballard, K. Balogh, K. Bender, B. Berger, M. Bogus, et al., Forensic validation of the SNPforID 52-plex assay, Forensic Science International: Genetics. 1 (2007) 186–190.

[13] C. Borsting, J.J. Sanchez, H.E. Hansen, A.J. Hansen, H.Q. Bruun, N. Morling, Performance of the SNPforID 52 SNP-plex assay in paternity testing, Forensic Science International: Genetics. 2 (2008) 292–300.

[14] A.J. Pakstis, W.C. Speed, R. Fang, F.C. Hyland, M.R. Furtado, J.R. Kidd, et al., SNPs for a universal individual identification panel, Human Genetics 127 (2010) 315–324.

[15] K.K. Kidd, J.R. Kidd, W.C. Speed, R. Fang, M.R. Furtado, F. Hyland, et al., Expanding data and resources for forensic use of SNPs in individual identification, Forensic Science International: Genetics. 6 (2012) 646–652.

[16] T.M. Karafet, F.L. Mendez, M.B. Meilerman, P.A. Underhill, S.L. Zegura, M.F. Hammer, New binary polymorphisms reshape and increase resolution of the human Y chromosomal haplogroup tree, Genome Res. 18 (2008) 830–838.

[17] Thermo Fisher Scientific, Precision ID SNP Panels with the HID Ion S5™/HID Ion GeneStudio™ S5 System Application Guide, (2023).

[18] Thermo Fisher Scientific, Technical Note: Maximize Forensic Genetic Analysis Potential Using an STR + SNP Combination NGS Workflow, (2019).

[19] S.M. Joo, Y. Kwon, M.H. Moon, K. Shin, Genetic investigation of 124 SNPs in a Myanmar population using the Precision ID Identity Panel and the Illumina MiSeq, Legal Medicine 63 (2023) 102256.

[20] C. Marshall, K. Sturk-Andreaggi, E.M. Gorden, J. Daniels-Higginbotham, S.G. Sanchez, Ž Bašić, et al., A Forensic Genomics Approach for the Identification of Sister Marija Crucifiksa Kozulić, Genes. 11 (2020) 938.

[21] A.C. Jäger, M.L. Alvarez, C.P. Davis, E. Guzmán, Y. Han, L. Way, et al., Developmental validation of the MiSeq FGx Forensic Genomics System for Targeted Next Generation Sequencing in Forensic DNA Casework and Database Laboratories, Forensic Science International: Genetics. 28 (2017) 52–70.

[22] C. Hollard, L. Ausset, Y. Chantrel, S. Jullien, M. Clot, M. Faivre, et al., Automation and developmental validation of the ForenSeq™ DNA Signature Preparation kit for high-throughput analysis in forensic laboratories, Forensic Science International: Genetics. 40 (2019) 37–45.

[23] R.E. Kieser, M.M. Buś, J.L. King, W.v.d. Vliet, J. Theelen, B. Budowle, Reverse Complement PCR: A novel one-step PCR system for typing highly degraded DNA for human identification, Forensic Science International: Genetics. 44 (2020) 102201.

[24] M.M. Bus, E.A. de Jong, J.L. King, W. van der Vliet, J. Theelen, B. Budowle, Reverse complement-PCR, an innovative and effective method for multiplexing forensically relevant single nucleotide polymorphism marker systems, BioTechniques. 71 (2021) 484–489.

[25] A.E. Woerner, J.L. King, B. Budowle, Fast STR allele identification with STRait Razor 3.0, Forensic Science International: Genetics. 30 (2017) 18–23.

[26] J.L. King, A.E. Woerner, S.N. Mandape, K.B. Kapema, R.S. Moura-Neto, R. Silva, et al., STRait Razor Online: An enhanced user interface to facilitate interpretation of MPS data, Forensic Science International: Genetics. 52 (2021) 102463.

[27] A. Staadig, J. Hedman, A. Tillmar, Applying Unique Molecular Indices with an Extensive All-in-One Forensic SNP Panel for Improved Genotype Accuracy and Sensitivity, Genes. 14 (2023) 818.

[28] R.C. Team, R: A language and environment for statistical computing, (2013).

[29] G.R. Abecasis, S.S. Cherny, W.O. Cookson, L.R. Cardon, Merlin--rapid analysis of dense genetic maps using sparse gene flow trees, Nature Genetics 30 (2002) 97–101.

[30] J. Snedecor, T. Fennell, S. Stadick, N. Homer, J. Antunes, K. Stephens, et al., Fast and accurate kinship estimation using sparse SNPs in relatively large database searches, Forensic Science International: Genetics 61 (2022) 102769.

[31] J. Watson, D. McNevin, K. Grisedale, M. Spiden, S. Seddon, J. Ward, Operationalisation of the ForenSeq® Kintelligence Kit for Australian unidentified and missing persons casework, Forensic Science International: Genetics 68 (2024) 102972.

[32] M.A. Peck, A.F. Koeppel, E.M. Gorden, J.L. Bouchet, M.C. Heaton, D.A. Russell, et al., Internal Validation of the ForenSeq Kintelligence Kit for Application to Forensic Genetic Genealogy, Forensic Genomics 2 (2022).

[33] S.M. Radecke, J. Antunes, J. Snedecor, C. Zlatkov, L. Devesse, G. Padmabandu, et al., Evaluation of a High-Throughput Dense Single-Nucleotide Polymorphism PCR Multiplex Next Generation Sequencing Application for Human Remains Identification, Forensic Genomics 3 (2023) 75–93.

[34] I.G.P. Consortium, A global reference for human genetic variation, Nature 526 (2015) 68.

[35] T.C. Matise, F. Chen, W. Chen, M. Francisco, M. Hansen, C. He, et al., A second-generation combined linkage–physical map of the human genome, Genome Res. 17 (2007) 1783–1786.

[36] Scientific Working Group on DNA Analysis Methods (SWGDAM), Recommendations of the SWGDAM Ad Hoc Working Group on Genotyping Results Reported as Likelihood Ratios, (2018).

[37] T.J. Parsons, R.M. Huel, Z. Bajunović, A. Rizvić, Large scale DNA identification: the ICMP experience, Forensic Science International: Genetics 38 (2019) 236–244.

